# Targeting Tumor Immunity: Statins as a Repurposed Strategy for Preventing Ovarian Cancer Advancement

**DOI:** 10.64898/2026.09.14.751179

**Authors:** Juliane M. Liberto, Sumeng Qi, Md Masud Rana, Ie-Ming Shih, Tian-Li Wang

## Abstract

High-grade serous ovarian carcinoma (HGSOC) remains the most lethal gynecologic malignancy, characterized by a high recurrence rate and poor long-term survival. Although daily statin use is associated with improved survival in ovarian cancer patients, the underlying mechanism of this benefit remains unclear, in part due to the use of non-clinical high doses and reliance on immunodeficient murine models in preclinical studies. Here, we analyzed data from the TriNetX US Collaborative Network to evaluate the association between statin use and survival in ovarian cancer patients, confirming a survival benefit in a large U.S. cohort. We next investigated the protective effects in a syngeneic ascites tumor model of ovarian cancer. Mice treated with clinically relevant doses of lipophilic atorvastatin (ATO) exhibited prolonged survival. Single-cell transcriptomic analyses further revealed enhanced CD8^+^ T-cell activation programs and pro-inflammatory macrophage remodeling. Flow cytometry identified increased M1-like macrophages and activated CD8^+^ T-cell populations. Cytokine profiling and transcriptomic analyses indicated that IL-15 signaling was significantly elevated following ATO treatment. Multiplex immunofluorescence analyses demonstrated enhanced tertiary lymphoid structure (TLS)-associated immune organization in metastatic tumor tissues after ATO treatment, with increased T- and B-cell markers. Ex vivo culture of primary T cells with ascites supernatant from ATO-treated mice resulted in enhanced T-cell activation and migratory capacity compared with supernatant from DMSO-treated mice. Collectively, our results suggest that daily statin use may remodel the body environment into a more immune-activated state, leading to a long-term beneficial effect on survival in cancer patients, as observed in large epidemiological studies.

## Introduction

High-grade serous ovarian carcinoma (HGSOC) is the most common and highly aggressive form of ovarian epithelial malignancy, with over half of all cases diagnosed at the advanced stages. Despite recent advances in surgical management, chemotherapy, and targeted therapies, long-term survival for HGSOC remains poor, with a 5-year survival rate below 50% (1,2). There is a pressing need for novel approaches to prevent and/or treat this disease. Developing new anti-cancer drugs is time-consuming and costly. Therefore, repurposing existing approved drugs for other medications as cancer therapeutics represents a promising alternative approach.

The potential role of statins as anti-cancer agents has been recognized for more than two decades, and statins have long been explored as potential anti-cancer therapeutics (3). Some early-phase clinical trials further investigated high-dose statin therapy in patients with advanced solid tumors, where statins showed manageable toxicity and preliminary anti-tumor activity (4). In addition, combination studies evaluating high-dose statins together with targeted therapies in advanced solid malignancies have further suggested the therapeutic potential of statin-based combination strategies (5).

Large-scale epidemiological studies in ovarian cancer have also supported the potential therapeutic benefit of statins in ovarian cancer patients (6–8). In a nationwide Finnish observational study of more than 10 000 patients with epithelial ovarian cancer (6), ever use of statins before ovarian cancer diagnosis was associated with approximately a 40% reduction in ovarian cancer mortality compared with never use. In addition, lipophilic statins, particularly atorvastatin and simvastatin, showed stronger protective associations than hydrophilic statins, suggesting potential differences in anti-tumor efficacy among statin subtypes (6). Similar findings were also reported in a large cohort study of Korean women aged 45-70 years, in which long-term statin use was associated with reduced mortality across multiple gynecologic malignancies, including ovarian cancer (8).

Several studies have reported the growth-suppressive effects of statins in murine tumor models. For example, preclinical studies in transgenic and xenograft mouse models have demonstrated that statins can suppress ovarian tumor growth and inhibit the formation of serous tubal intraepithelial carcinomas (STICs), a precursor lesion of HGSOC (9). However, a clear mechanism by which statins restrain in vivo cancer progression, and whether it can be through modulation of the tumor microenvironment (TME), remains unclear.

Much of the prior research relied on immunodeficient murine models for in vivo studies, with most experiments using doses substantially higher than clinically relevant doses for cardiovascular diseases (9–11), thereby limiting their translational applicability. This study was thereby designed to assess potential anti-tumor modulatory effects in statin daily users. We used an immunocompetent murine model of ovarian cancer, and administered a widely prescribed lipophilic statin, atorvastatin, at a clinically relevant dose, and its effects on survival and immune landscape remodeling were evaluated in detail using integrated approaches.

## Materials and Methods

### Population-Based Survival Analysis

A retrospective cohort analysis was conducted using the TriNetX US Collaborative Network. Ovarian cancer patients were identified using the ICD-10 code C56 (malignant neoplasm of the ovary). Patients with documented use of HMG-CoA reductase inhibitors (statins) were assigned to the statin-user cohort, whereas those without statin exposure were assigned to the non-user cohort. Propensity score matching was performed on the TriNetX platform using all listed characteristics. After matching, 40 434 patients remained in each cohort. Mortality was the outcome of interest. The index event was defined as the first recorded diagnosis of ovarian cancer. Outcomes occurring from 1 day to 1 825 days after the index event were included in the analysis. Mortality risk was compared between cohorts using risk analysis. Overall survival was evaluated using Kaplan-Meier survival analysis, and differences between survival curves were assessed using the log-rank test. Hazard ratios (HRs) with 95% confidence intervals (CIs) were calculated to estimate the relative risk of death between cohorts.

### Syngeneic HGSOC Mouse Model and Atorvastatin Treatment

Wild-type female C57BL/6J mice (6-8 weeks old) were used for in vivo experiments. Mice were randomly assigned to receive daily intraperitoneal treatment with atorvastatin (ATO, 5 mg/kg/day) or vehicle control (DMSO, <2% v/v) for two weeks before tumor implantation. To establish a syngeneic high-grade serous ovarian cancer (HGSOC) mouse model, we injected mice intraperitoneally with 5 × 10⁵ ID8/Brca1-null cells (12). The same treatment was then continued until the experimental endpoint at 4 weeks post-tumor injection, corresponding to a total treatment duration of 6 weeks. Body weight was monitored longitudinally throughout tumor progression.

For endpoint analyses, mice were euthanized at designated experimental time points. Ascites fluid was collected, filtered through a 70-μm cell strainer, and centrifuged to separate cellular and supernatant fractions for downstream analyses. Cell pellets were subjected to red blood cell lysis before further processing. Peripheral blood was collected in EDTA-containing tubes and centrifuged to isolate plasma. Peritoneal tissues containing metastatic tumor lesions were harvested, fixed in formalin, embedded in paraffin, sectioned, and processed for histological and multiplex immunofluorescence analyses.

For survival studies, mice were monitored until reaching institutional humane endpoint criteria. Survival analysis was performed on the integrated data of two independent experiments. Ascites volume was measured at the time of necropsy.

All animal experiments were performed in accordance with institutional guidelines and approved by the Institutional Animal Care and Use Committee (IACUC) of Johns Hopkins University.

### 10x Genomics Fixed RNA Profiling and Library Preparation

CD45^+^ live immune cells were isolated from ascites samples collected from ATO- and DMSO-treated ovarian tumor-bearing C57BL/6J mice by fluorescence-activated cell sorting (FACS). Approximately 1 × 10⁶ CD45^+^ live cells were fixed using the Chromium Fixed RNA Profiling protocol (10x Genomics) according to the manufacturer’s instructions. Briefly, sorted cells were pelleted, washed, resuspended in fixation, incubated overnight, and quenched after approximately 20 hours by quenching buffer. Cell concentrations were determined based on the trypan blue exclusion method, which was performed on a Countess III cell counter. Fixed cells were stored at −80°C in enhancer buffer supplemented with 10% glycerol before library preparation.

Single-cell RNA sequencing libraries were prepared using the Chromium Fixed RNA Profiling workflow for multiplexed samples (10x Genomics) according to the manufacturer’s protocol. Fixed immune cells were thawed and hybridized with mouse whole transcriptome analysis (WTA) probes. Following probe hybridization, samples were pooled, washed, filtered, and processed for GEM generation and barcoding using the Chromium Next GEM Chip Q on the Chromium X/iX instrument. GEMs were recovered, pre-amplified, indexed, purified by size selection, and assessed for library quality before sequencing.

## Results

### Statin use is associated with improved survival among ovarian cancer patients and prolongs survival in a syngeneic HGSOC model

To investigate the clinical relevance of statin use in ovarian cancer, we analyzed a large real-world patient cohort from the TriNetX database. After propensity score matching, 40 434 patients remained in each cohort. Statin users exhibited a significantly lower mortality risk compared with non-users (20.9% vs. 23.2%) (p < 0.001, **Figure 1A**), corresponding to a risk ratio of 0.90 and an odds ratio of 0.87 **(Figure 1A)**. Kaplan-Meier survival analysis further demonstrated significantly improved overall survival among statin users compared with non-users (p < 0.001, log-rank test, **Figure 1B**). These findings suggest that statin use is associated with improved survival outcomes in ovarian cancer patients.

**Figure 1.**
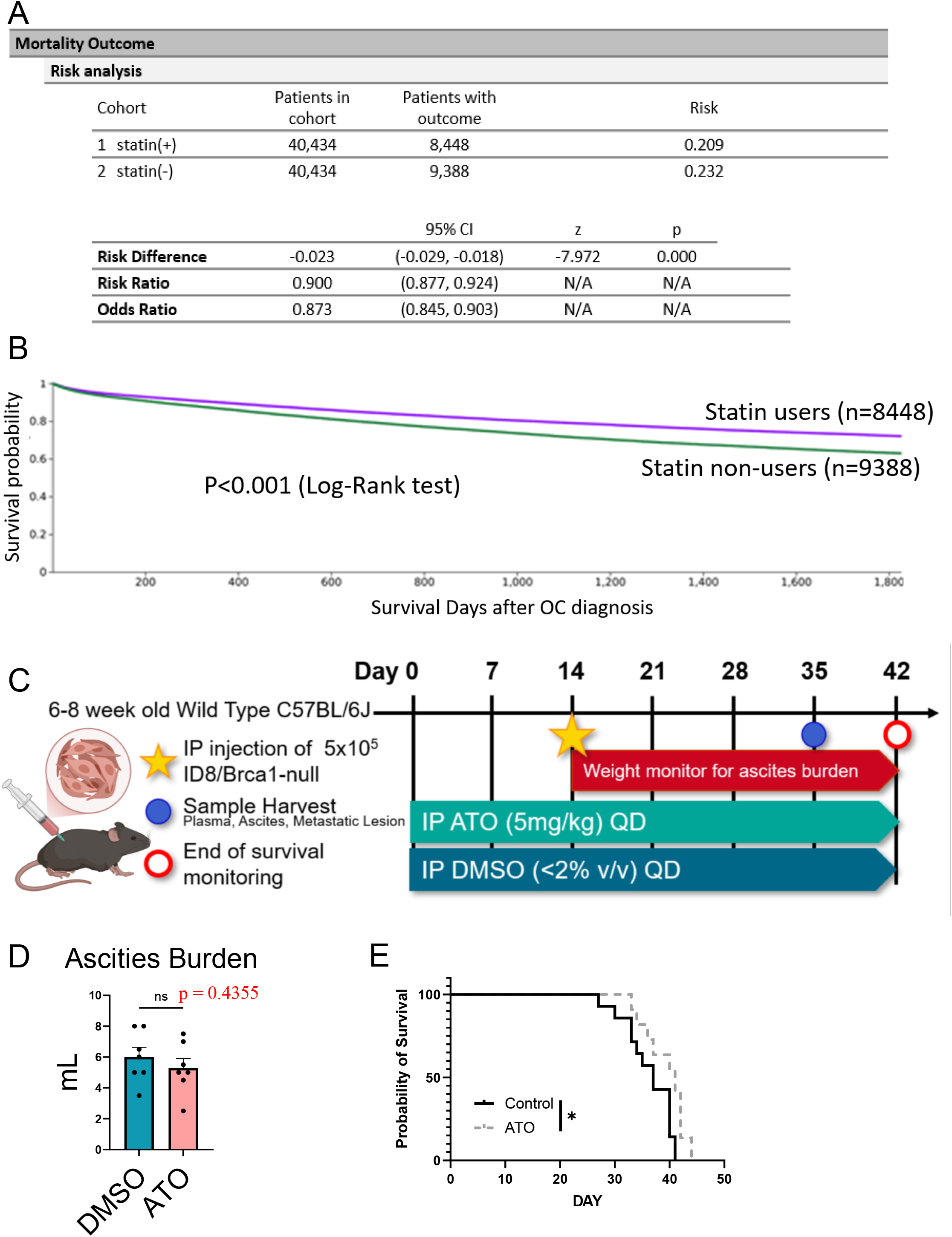
Statin use is associated with prolonged survival in ovarian cancer patients. **(A)** Risk analysis comparing mortality outcomes between statin users (1) and non-users (2) in ovarian cancer patients after propensity score matching. The table summarizes the number of patients, mortality risk, risk difference, risk ratio, and odds ratio between the two cohorts. **(B)** Kaplan-Meier overall survival curves of propensity score-matched ovarian cancer patients with or without statin use. Survival probability was analyzed over a 5-year follow-up period after ovarian cancer diagnosis. Statistical significance was determined using the log-rank test. **(C)** Schematic of the HGSOC tumor-bearing mice experimental design. Female C57BL/6J mice aged 6-8 weeks were pretreated with daily intraperitoneal atorvastatin (ATO, 5 mg/kg) or vehicle control (DMSO, <2% v/v) for 2 weeks before intraperitoneal injection of ID8/Brca1-null cells. The same daily treatment was then continued after tumor cell injection for a total treatment duration of 6 weeks. Body weight was monitored longitudinally, and mice were allocated for endpoint ascites and tissue collection or survival analysis. **(D)** Bar chart showing the volume of ascites collected at week 5 after the start of ATO treatment. Data are presented as mean ± SEM. Statistical analysis was performed using an unpaired two-tailed Student’s t-test; ns, not significant. **(E)** Kaplan-Meier survival analysis of ATO- and DMSO-treated mice. Statistical significance was determined using the log-rank (Mantel-Cox) test. Data are presented as mean ± SEM. *P < 0.05, **P < 0.01, ***P < 0.001, ****P < 0.0001; ns, not significant.

To investigate the effect of statin intake at clinical doses prescribed for cardiovascular disease on preventing or delaying cancer progression, we established a syngeneic high-grade serous ovarian carcinoma (HGSOC) mouse model by intraperitoneal injection of ID8/Brca1-null cells into C57BL/6J mice. ID8/Brca1-null is an aggressive ovarian tumor model that readily forms ascites tumors within two weeks after tumor cell injection. Mice received daily intraperitoneal treatment with atorvastatin (ATO, 5 mg/kg) or vehicle control for a total of six weeks, as depicted in **Figure 1C**.

Body weight was monitored longitudinally throughout tumor progression, and no significant differences were observed between DMSO- and ATO-treated mice during the experimental period **(Supplementary Figure 1A)**. Additionally, ascites volume measured at the endpoint was not significantly altered between ATO-treated and DMSO-treated groups **(Figure 1D)**. Gross necropsy examination and H&E staining of metastatic tumor tissues showed comparable overall tumor morphology between groups **(Supplementary Figure 1B)**. These findings suggest that ATO treatment at this clinically prescribed dose for lowering blood cholesterol did not substantially alter overall tumor burden in this model. Despite the absence of a significant reduction in tumor burden, ATO-treated mice exhibited prolonged overall survival compared with DMSO-treated mice as demonstrated by the Kaplan-Meier survival analysis **(Figure 1E)**. Most mice in the DMSO-treated group died between 30 and 40 days after tumor implantation, whereas ATO treatment extended survival.

### Statin treatment remodels T-cell landscapes and cytokine-response programs in the ascites tumor microenvironment

To further characterize immune changes associated with ATO treatment, CD45^+^ immune cells were isolated from tumor ascites of DMSO- and ATO-treated HGSOC tumor-bearing mice and analyzed by 10x Flex single-cell RNA sequencing (scRNA-seq) **(Figure 2A)**. Unsupervised clustering identified multiple immune and stromal cell populations in tumor ascites, including T cells, B cells, dendritic cells, macrophages, monocytes, granulocytes, NK cells, fibroblasts, and microglia, which are presented as a UMAP in **Figure 2B**. A three-dimensional UMAP of these immune cell clusters is shown in the **Supplementary File**.

**Figure 2.**
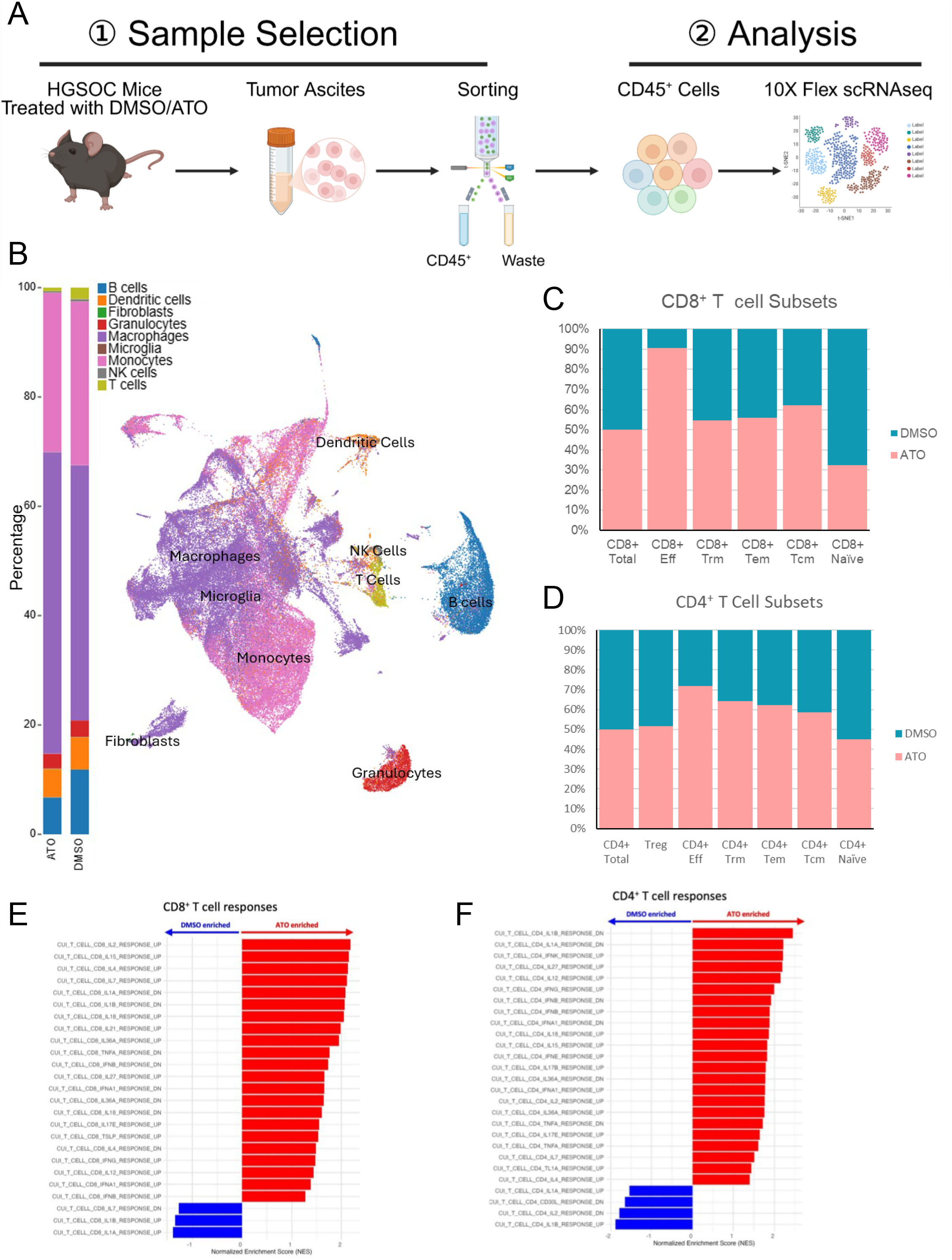
scRNA-seq analysis reveals T-cell transcriptional remodeling in statin-treated HGSOC tumors. **(A)** Schematic of the single-cell RNA sequencing workflow. CD45+ immune cells were isolated from tumor ascites collected from DMSO- and ATO-treated HGSOC tumor-bearing mice, followed by 10x Genomics Flex scRNA-seq analysis. Downstream analyses focused on T-cell and macrophage populations. **(B)** UMAP visualization of immune cell populations identified by scRNA-seq analysis, including T cells, B cells, dendritic cells, macrophages, monocytes, granulocytes, NK cells, fibroblasts, and microglia. **(C-D)** Relative frequencies of CD8⁺ T-cell subsets **(C)** and CD4⁺ T-cell subsets **(D)** in DMSO- and ATO-treated HGSOC tumor-bearing mice. **(E-F)** NES analysis of cytokine response programs in CD8⁺ T cells **(E)** and CD4⁺ T cells **(F)** from ATO-treated versus DMSO-treated HGSOC tumor-bearing mice. Positive NES values (red) indicate pathways enriched in ATO-treated T-cell populations, while negative NES values (blue) indicate pathways enriched in DMSO-treated T-cell populations.

T-cell populations were subsequently selected for focused transcriptional analysis **(Supplementary Figure 2A)**. Differential gene expression analysis revealed broad transcriptional remodeling in T cells following ATO treatment. Significantly upregulated genes included those associated with chemokine and cell migration, such as *Cxcl1*, *Ccr9*, *Cxcr3*, and *Ccl5*, as well as those involved in co-stimulation and activation, like *Cd28*, *Cd44*, *Cd48*, *Il2rb*, and *Cd160* **(Supplementary Figure 2A, Supplementary Table)**. T effector-associated genes, including *Gzmb*, *Gzma*, and *Nkg7*, were also upregulated **(Supplementary Figure 2A, Supplementary Table)**. Additionally, transcription factors linked to T-cell activation and differentiation, such as *Tbx21* and *Gata3*, were upregulated following ATO treatment **(Supplementary Figure 2A, Supplementary Table)**. In contrast, regulatory-associated genes, such as *Relb* and *Clec4a3*, were downregulated following ATO treatment **(Supplementary Figure 2A, Supplementary Table)**. Functional grouping analysis based on the top upregulated and downregulated genes identified from the differential expression analysis further demonstrated increased expression of activation/co-stimulation and cytoskeleton/migration genes in T-cell populations from ATO-treated mice **(Supplementary Figure 2B)**. In contrast, transcriptional programs associated with regulatory T cells showed reduced expression following ATO treatment **(Supplementary Figure 2B)**. Hallmark pathway enrichment analysis based on these top differentially expressed genes further identified enrichment of pathways associated with IFN-γ response, IL2-STAT5 signaling, mTOR signaling, and glycolysis in T cells from ATO-treated mice **(Supplementary Figure 2C)**. Enrichment of MYC targets and G2M checkpoint pathways further suggested increased proliferative and functional activation programs in T cells from ATO-treated mice. Conversely, TNFα signaling via NFκB and hypoxia-associated pathways were negatively enriched **(Supplementary Figure 2C)**.

To further characterize the T-cell landscape associated with statin treatment in our model, T-cell populations identified by scRNA-seq were selected for subclustering analysis **(Figure 2C-D)**. T-cell subsets were annotated based on representative marker-gene expression patterns and transcriptional features, including naïve, effector (Teff), effector memory (Tem), central memory (Tcm), tissue-resident memory (Trm), and regulatory T-cell states (Treg) **(Supplementary Figure 3A-B)**. Relative-frequency analysis of CD8^+^ T-cell subsets showed a reduction in naïve CD8^+^ T cells and enrichment of effector-associated CD8^+^ T-cell populations following ATO treatment **(Figure 2C)**. CD4^+^ T-cell populations showed moderate compositional changes between ATO-treated and control groups, with increases across effector and memory subsets **(Figure 2D)**.

We next examined lineage-specific transcriptional changes in CD4^+^ and CD8^+^ T cells. Pathway enrichment analysis demonstrated broad enrichment of cytokine-response-associated transcriptional programs in both CD8^+^ and CD4^+^ T-cell populations from ATO-treated mice **(Figure 2E-F)**. In CD8^+^ T cells, pathways associated with IL-2, IL-15, IFNγ, IFNα, IL-18, IL-21, and IL-27 responses were positively enriched following ATO treatment **(Figure 2E)**. Similarly, CD4^+^ T cells exhibited enrichment of IFNγ-, IL-15-, IL-18-, IL-27-, and TNFα-associated response programs **(Figure 2F)**. These findings suggest enhanced cytokine-responsive and activation-associated transcriptional states in both CD8^+^ and CD4^+^ T-cell populations following statin treatment.

### Statin treatment promotes inflammatory remodeling of macrophage populations in the ascites tumor microenvironment

To further characterize macrophage-associated immune remodeling following statin treatment, macrophage populations identified by scRNA-seq analysis were selected for focused transcriptional analysis. Broad transcriptional changes in macrophages from ATO-treated mice compared to the DMSO group were revealed by differential gene expression analysis **(Figure 3A)**. Upregulated genes included inflammatory- and immune-response-associated genes, such as *Plk2*, *Rasgef1b*, *Cxcl2*, and *Bhlhe40*, whereas anti-inflammatory or regulatory-associated genes like *Tgfb1* and *F13a1* were downregulated following ATO treatment **(Figure 3B)**. Consistently, analysis of macrophage polarization markers demonstrated enrichment of M1-like signatures and a relative reduction of M2-like signatures following ATO treatment **(Figure 3C)**. Pathway enrichment analysis using the macrophage-specific immune gene set (13) further demonstrated enrichment of inflammatory-response-associated pathways in macrophages from ATO-treated mice, including IL-15, TNFα, IFNγ, and NFκB-associated response programs **(Figure 3D)**. These findings indicate that ATO treatment induces inflammatory and M1-like transcriptional remodeling in tumor-associated macrophages.

**Figure 3.**
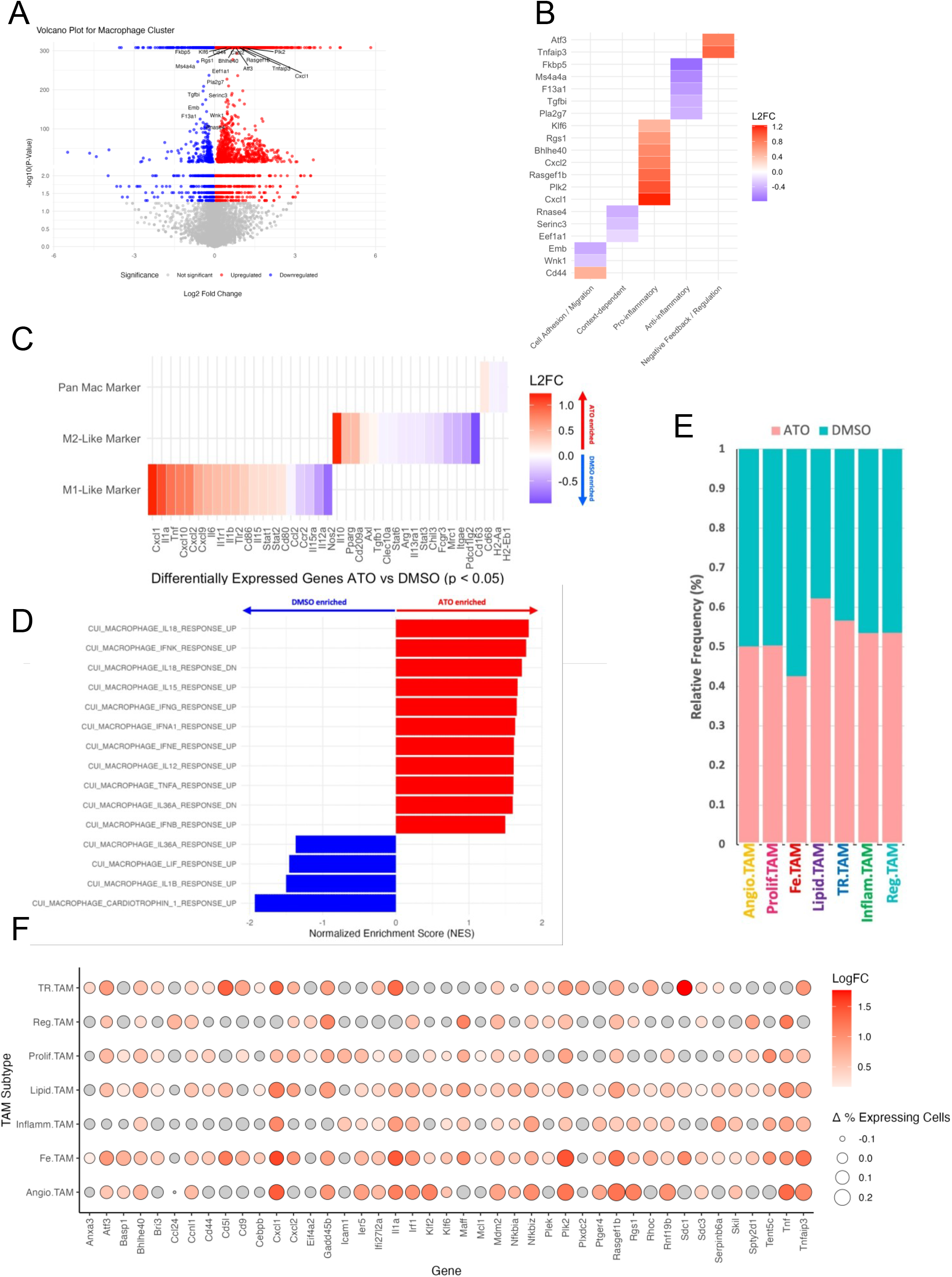
Statin treatment remodels macrophage states toward inflammatory and immune-supportive programs in HGSOC tumor-bearing mice. **(A)** Volcano plot showing differentially expressed genes in macrophage clusters from ATO-treated versus DMSO-treated mice. **(B)** Functional grouping analysis of differentially expressed genes associated with macrophage migration, cytoskeleton regulation, pro-inflammatory signaling, anti-inflammatory signaling, and negative feedback/regulatory programs following ATO treatment. **(C)** Heatmap of top differentially expressed genes associated with pan-macrophage, M1-like macrophage, and M2-like macrophage transcriptional programs in ATO-treated versus DMSO-treated mice. **(D)** Macrophage pathway enrichment analysis showing NES values for inflammatory and cytokine-response-associated transcriptional programs in macrophage populations from ATO-treated versus DMSO-treated mice. Positive NES values (red) indicate pathways enriched in ATO-treated macrophages, while negative NES values (blue) indicate pathways enriched in DMSO-treated macrophages. **(E)** Relative frequency of TAM-associated macrophage states in DMSO- and ATO-treated macrophage populations. **(F)** Dot plot showing subtype-specific expression changes of representative ATO-upregulated differentially expressed genes across TAM subtypes. Dot color indicates log fold change in ATO-treated versus DMSO-treated macrophages, with grey indicating no significant change within that TAM subtype. Dot size represents the change in the fraction of cells expressing each gene between the ATO and DMSO groups (% of ATO cells expressing gene − % DMSO cells expressing gene).

To further characterize macrophage heterogeneity in the ascites microenvironment, macrophage populations were annotated for TAM-associated transcriptional states, identifying seven tumor-associated macrophage states, including iron-handling TAMs (Fe.TAM), lipid-associated TAMs (Lipid.TAM), tissue-resident TAMs (TR.TAM), proliferative TAMs (Prolif.TAM), angiogenic TAMs (Angio.TAM), regulatory TAMs (Reg.TAM), and inflammatory TAMs (Inflam.TAM) **(Supplementary Figure 4A)**. Representative marker genes used for TAM-state annotation are listed in **Supplementary Figure 4A**, and the expression distributions of selected representative markers across the macrophage UMAP embedding are shown in **Supplementary Figure 4B**. Relative frequency analysis showed broadly similar distributions of most TAM states between treatment groups, with lipid-associated TAMs being relatively enriched in the ATO-treated mice **(Figure 3E)**. Dot plot analysis further highlighted heterogeneous gene-level responses to ATO treatment across TAM states **(Figure 3F)**. Fe.TAM and Lipid.TAM displayed broad transcriptional changes, with increased expression of several ATO-responsive genes, including *Cxcl1*, *Il1a*, *Plk2*, *Rasgef1b*, and *Tnfaip3*. Angio.TAM showed prominent induction of selected genes, including *Cxcl1*, *Il1a*, *Plk2*, *Rasgef1b*, *Rgs1*, and *Tnfaip3*, whereas TR.TAM exhibited strong increases in genes such as *Cd5l*, *Cxcl1*, *Il1a*, *Sdc1*, *Plk2*, and *Tnfaip3*. In contrast, Inflam.TAM, Reg.TAM, and Prolif.TAM exhibited more restricted or gene-specific changes, with fewer genes showing strong induction or increased fractions of expressing cells.

### Statin treatment promotes IL-15-associated immune remodeling and TLS activation in the ovarian cancer microenvironment

To investigate whether statin treatment alters mouse immune cell composition, flow cytometric analyses were performed on immune populations isolated from the spleens of HGSOC tumor-bearing mice treated with DMSO or ATO **(Figure 4A-B, Supplementary Figure 5A, 6A-B)**. Analysis of myeloid cell populations showed no significant differences in the overall frequencies of conventional dendritic cells (cDCs), total myeloid-derived suppressor cells (MDSCs), granulocytic MDSCs (G-MDSCs), monocytic MDSCs (M-MDSCs), M2-like macrophages, or total macrophages between treatment groups **(Supplementary Figure 6A)**. However, ATO-treated mice showed a significant increase in M1-like macrophages compared to the DMSO group **(Figure 4A; p < 0.05, Student’s t-test)**. Flow cytometric analysis of T-cell populations demonstrated no major differences in total T cells, CD4^+^ T cells, or CD8^+^ T cells between ATO- and DMSO-treated groups **(Supplementary Figure 6B)**. In contrast, activated CD8^+^ T cells showed a marginal increase in the ATO-treated group compared to the DMSO-treated group (**Figure 4B**, p = 0.0719, Student’s t-test).

**Figure 4.**
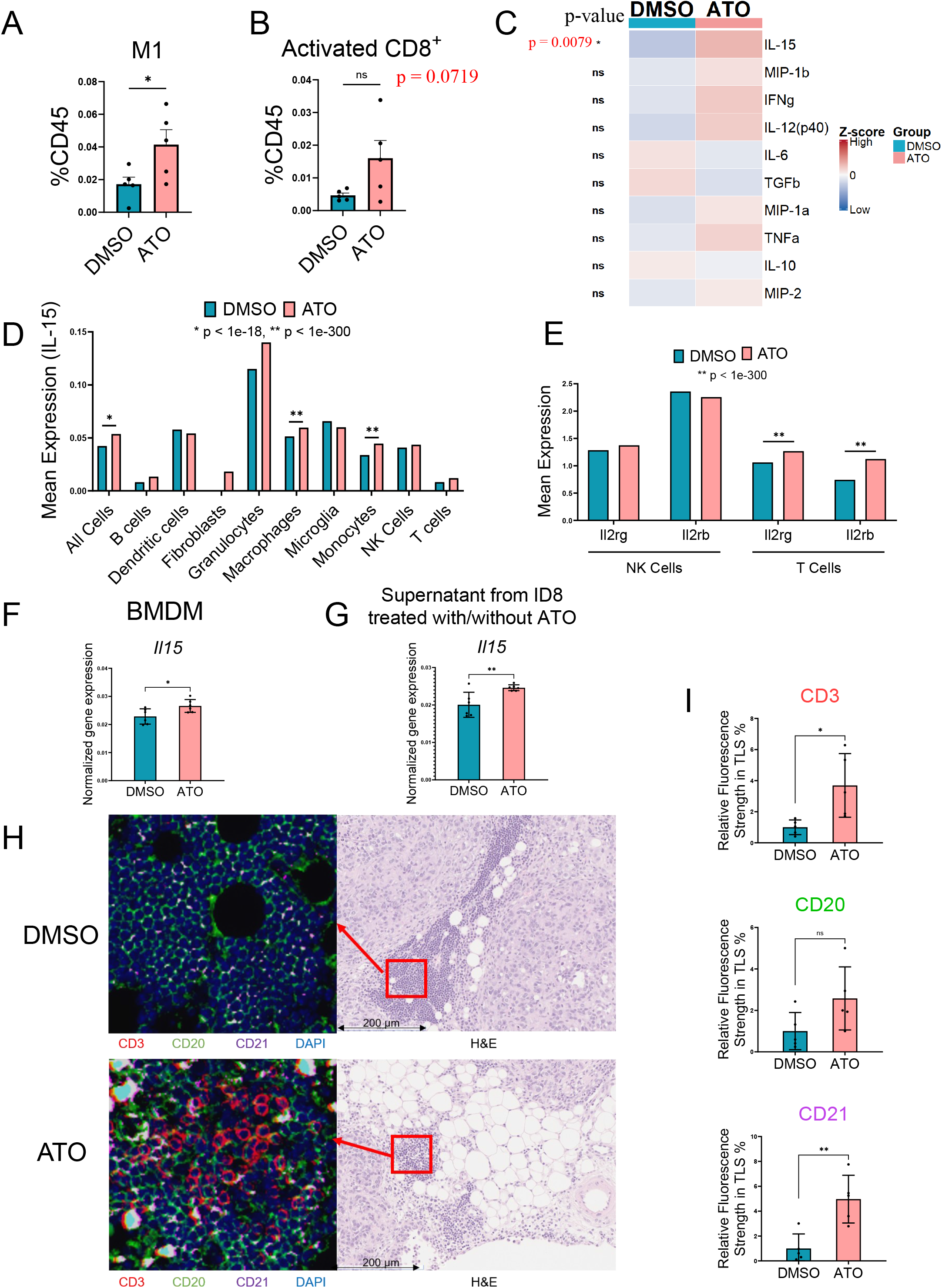
Statin treatment enhances IL-15-associated immune remodeling and TLS activation in the ovarian cancer microenvironment. **(A-B)** Flow cytometric quantification of selected immune cell populations in spleens collected from DMSO- and ATO-treated HGSOC tumor-bearing mice, including **(A)** M1-like macrophages and **(B)** activated CD8+ T cells. **(C)** Multiplex cytokine analysis of ascites supernatants showing IL-15 levels in DMSO- and ATO-treated HGSOC tumor-bearing mice. **(D)** Mean expression levels of IL-15 across major immune and stromal cell populations identified by scRNA-seq analysis from ATO- and DMSO-treated HGSOC tumor-bearing mice. **(E)** Mean expression levels of *Il2rg* and *Il2rb* in NK-cell and T-cell populations identified by scRNA-seq analysis from ATO- and DMSO-treated HGSOC tumor-bearing mice. **(F-G)** Quantitative PCR analysis of *Il15* expression in **(F)** BMDMs treated ex vivo with vehicle control or 2 μM ATO for 4 hours, or **(G)** BMDMs cultured with supernatants collected from ID8 ovarian cancer cells treated with vehicle control or 2 μM ATO for 24 hours. **(H)** Representative multiplex immunofluorescence staining and corresponding H&E images of metastatic peritoneal tumor sections collected from DMSO- and ATO-treated HGSOC tumor-bearing mice. Regions enriched for lymphocyte aggregates and morphologically consistent with TLS-like areas were selected for analysis. Multiplex immunofluorescence staining was performed for CD3 (red, T cells), CD20 (green, B cells), CD21 (purple, mature B cells), and DAPI (blue). Red boxes in H&E images indicate the corresponding regions shown in multiplex immunofluorescence images. Scale bar, 200 μm. **(I)** Quantification of relative fluorescence intensity of CD3, CD20, and CD21 signals within TLS-like regions. Statistical analysis was performed as follows: **(D-E)** Two-sample t-test with Welch’s correction was performed on log-normalized expression values in Cirrocumulus, followed by Benjamini-Hochberg correction for multiple testing. Adjusted P values < 0.05 were considered significant, and the significance thresholds are indicated separately as *P < 1 × 10^-18^ and **P < 1 × 10^-300^; **(A-C, F-G, I)**: Unpaired two-tailed Student’s t-test was performed. Data are presented as mean ± SEM. For these panels, *P < 0.05, **P < 0.01, ***P < 0.001, ****P < 0.0001; ns, not significant.

To further investigate systemic and local cytokine responses associated with ATO treatment, we performed multiplex cytokine analyses of plasma and ascites supernatants collected from treated mice **(Figure 4C, Supplementary Figure 6C)**. Most cytokines analyzed did not show significant differences between ATO and DMSO treatment groups. However, IL-15 levels were significantly elevated in ascites supernatant of the ATO-treated mice **(Figure 4C; p < 0.01, Student’s t-test)**. Additionally, IL-15 levels in the plasma of ATO-treated mice showed a marginal increase **(Supplementary Figure 6C, p = 0.053, Student’s t-test)**. Together, these findings suggest systemic immune activation-associated changes following statin treatment, accompanied by increased IL-15-associated cytokine responses.

To further investigate IL-15-associated signaling within the ATO-treated tumor microenvironment, scRNA-seq data were analyzed for *Il15* expression among immune cell populations **(Figure 4D)**. Increased Il15 expression following ATO treatment was primarily observed in myeloid cell populations, including macrophages and monocytes **(Figure 4D)**. Because IL-15 signaling utilizes IL2RB and IL2RG receptor subunits shared between T cells and NK cells, expressions of *Il2rb* and *Il2rg* were further examined in these lymphocyte populations. T cells from ATO-treated mice exhibited increased expression of both *Il2rb* and *Il2rg* compared with the DMSO group, whereas NK cells showed relatively limited changes **(Figure 4E)**. To further determine whether ATO directly regulates macrophage-associated IL-15 expression, bone marrow-derived macrophages (BMDMs) were cultured ex vivo and treated with ATO (14,15). qRT-PCR analysis showed increased *Il15* expression in BMDMs following direct ATO treatment **(Figure 4F)**. Similarly, macrophages cultured with supernatants from ATO-treated ID8 ovarian cancer cells also showed increased *Il15* expression compared with controls **(Figure 4G)**. Together, these findings suggest that ATO treatment promotes IL-15-associated immune signaling within the ovarian cancer tumor microenvironment.

Ovarian cancer can present in both ascites and solid tumor forms, and the ID8 model simulates not only ascites tumor generation but also tumor implantation (or metastasis) into peritoneal tissues. We next evaluated whether ATO treatment alters the peritoneal tumor tissue microenvironment of the ID8 tumor-bearing mice. Pathological evaluation of H&E-stained peritoneal tissue sections identified lymphocyte-rich regions morphologically consistent with tertiary lymphoid structures (TLSs), including dense clusters of small, darkly stained nuclei, indicative of localized immune cell aggregation **(Supplementary Figure 7A)**. Multiplex IF staining was performed to evaluate T-cell- and B-cell-associated markers, including CD3, CD20, and CD21 **(Figure 4H)**. Quantification of fluorescence intensity from the immunostaining demonstrated increased CD3^+^ T-cell signals in TLS-like regions of ATO-treated mice compared with the DMSO group **(Figure 4I)**. CD20, a pan-B-cell marker, showed a modest increase following ATO treatment, but the result was not statistically significant. In contrast, expression of CD21, a marker for TLS maturation, was significantly increased in the ATO-treated mice **(Figure 4I)**.

### Ascites supernatant of the statin-treated mice modifies T-cell activation states in ex vivo cultures

To investigate whether the statin-remodeled tumor ascites microenvironment directly affects T-cell activation, we established an ex vivo primary T-cell culture system and incubated it with cell-free ascites supernatants purified from the DMSO-or ATO-treated ID8/Brca1-null tumor-bearing mice (experimental schema depicted in **Figure 5A**). Primary CD3^+^ T cells were isolated from the spleens of C57BL/6J mice, activated with anti-CD3/CD28 and IL-2 cytokine, and subsequently cultured with tumor ascites supernatants for 24 hours. qRT-PCR analysis showed that after incubation of activated primary CD3^+^ T cells with ascites supernatant from the ATO-treated mice, *Il2* expression was increased compared to the DMSO group **(Figure 5B)**. Expression of *Ltb*, a cytokine associated with TLS formation, was also elevated. These findings indicate enhanced T-cell activation under the ATO-treated tumor microenvironment. Meanwhile, increased expression of *Pdcd1* was also observed in T cells exposed to ascites supernatants from ATO-treated mice **(Figure 5B)**.

**Figure 5.**
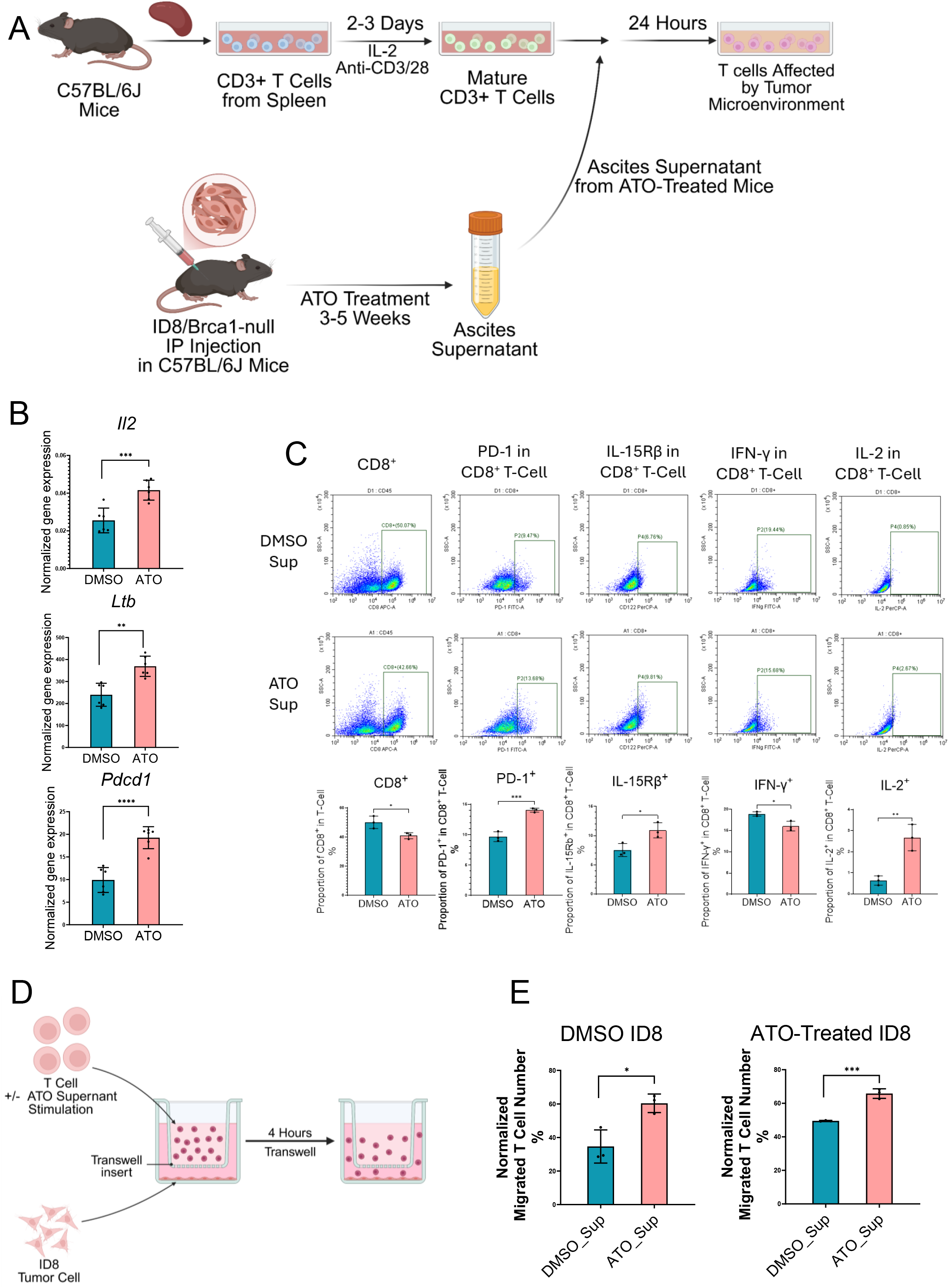
Tumor ascites from atorvastatin-treated mice alters T-cell phenotypes and functional responses ex vivo. **(A)** Schematic of the ex vivo primary T-cell experimental design. CD3^+^ T cells were isolated from spleens of C57BL/6J mice and activated with IL-2 and anti-CD3/CD28 stimulation for 2-3 days. Activated T cells were subsequently cultured for 24 hours in the presence of ascites supernatants collected from DMSO-or ATO-treated HGSOC tumor-bearing mice. **(B)** Quantitative PCR analysis of *Il2*, *Ltb*, and *Pdcd1* expression in ex vivo T cells cultured with DMSO-or ATO-treated ascites supernatants. **(C)** Flow cytometric analysis of ex vivo CD8⁺ T cells cultured with DMSO- or ATO-treated ascites supernatants. CD8⁺ T cells were analyzed for the expression of PD-1, IL-15Rβ, IFN-γ, and IL-2 after culture with ascites supernatants from DMSO-or ATO-treated mice. Representative flow cytometry plots and corresponding quantifications are shown. **(D)** Schematic of the transwell migration assay using ex vivo T cells cultured with DMSO- or ATO-treated ascites supernatants. **(E)** Quantification of migrated T cells following transwell assays toward DMSO- or ATO-treated ID8 tumor cells. Migrated T cells were collected and quantified by flow cytometry. Statistical analysis was performed using an unpaired two-tailed Student’s t-test. Data are presented as mean ± SEM. *P < 0.05, **P < 0.01, ***P < 0.001, ****P < 0.0001; ns, not significant.

To further characterize T-cell phenotypic changes, flow cytometric analyses were performed separately on CD4^+^ and CD8^+^ T-cell populations **(Figure 5C, Supplementary Figure 8A)**. In CD4^+^ T cells, IL-2 expression was increased following exposure to ascites supernatants of the ATO-treated mice, while IFN-γ expression showed a modest upward **trend (Supplementary Figure 8A, p = 0.0413, Student’s t-test)**. In contrast, PD-1 and IL-15Rβ expression did not show significant changes in the CD4^+^ T-cell population. In CD8^+^ T cells, IL-2 and PD-1 expressions were increased following exposure to ascites supernatants of the ATO-treated mice, whereas IFN-γ expression was decreased compared to the DMSO group **(Figure 5C)**. Additionally, IL-15Rβ expression was elevated in CD8^+^ T cells following exposure to ATO-conditioned ascites supernatants. Since IL-15 cytokine levels were found to be elevated in ascites **(Figure 4C)**, the results together indicate a strong activation of the IL-15/IL-15Rβ signaling axis in the ATO-treated ascites tumor microenvironment.

To investigate whether the ATO-conditioned tumor microenvironment modifies T-cell migratory capacity, a transwell migration assay was performed on ex vivo T cells cultured with ascites supernatant (experimental schema shown in **Figure 5D**). Primary T cells exposed to ascites supernatants of DMSO-or ATO-treated mice were seeded into the upper transwell inserts. Meanwhile, ID8 tumor cells were seeded in the lower wells. T cells that migrated to the lower chamber were assessed by flow cytometry. As shown in **Figure 5E**, incubating T cells with ascites supernatants from ATO-treated mice increased their migratory capacity, regardless of whether the tumor cells were present in the lower well. These findings indicate that exposure to the ATO-conditioned ascites microenvironment promotes T-cell migratory capacity.

## Discussion

Previous epidemiological studies have indicated that statin use is associated with better survival outcomes in ovarian cancer patients after diagnosis (6–8). Preclinical studies in cancer cell lines and animal models have further supported this observation. However, most preclinical studies applied supraphysiologic doses to cancer cell lines, and in vivo studies often relied on immunodeficient murine models, which collectively limited the context relevance and direct clinical applicability (9–11).

Using large-scale ovarian cancer data cohorts from US cooperative groups, we discovered that statin users have better overall survival than non-users **(Figure 1A-B)**. To elucidate potential mechanisms, we investigated the effects of a daily clinical dose of atorvastatin (ATO) in a syngeneic ascites HGSOC murine model, which recapitulates ascites formation and peritoneal metastasis of advanced ovarian cancer. We found that even at this relatively low dose used for cardiovascular disease prevention, the treatment was associated with a statistically significant prolongation of survival and broad remodeling of the tumor microenvironment (TME). Nevertheless, at the current dose, ATO did not significantly suppress tumor burden despite improving survival. One possibility is that the low daily statin dose was insufficient to directly suppress tumor growth compared with cytotoxic chemotherapeutic drugs. Despite limited direct effects on tumor cells, significant remodeling of the host tissue environment was observed. These included an enhanced CD8^+^ adaptive immune program in ascites and an increased presence of lymphoid structures in peritoneal tissues. The findings indicate that the host body, including the immune system, can respond to inhibition of the mevalonate pathway even at low doses.

Mechanistically, tumor cells and immune cells likely have distinct metabolic preferences and dependencies, thereby presenting differential sensitivities to statin treatment. To support fast growth and environmental challenges, cancer cells rely heavily on multiple metabolic pathways, including glucose and glutamine metabolism, which may partially buffer the effects of mevalonate pathway inhibition. In contrast, normal cells, including specific immune cells, may be more sensitive to metabolic changes induced by low-dose statins and elicit remodeling of the tissue environment. Such a feature could offer translational advantages, as low-dose statin use could potentiate anti-tumor immunity in a less toxic and more clinically feasible manner.

Single-cell transcriptomic analyses suggest that statin treatment reshapes the ascites tumor immune microenvironment toward a more inflammatory, adaptive, and immune-active state. In particular, statin treatment appeared to favor adaptive immune populations, including increased overall T-cell and B-cell abundance. Pathway analyses further suggested substantial remodeling of metabolic pathways in T-cell programs after statin treatment. Consistent with these findings, increased effector and memory-associated T cells **(Figure 2C-D)** further support more functionally adaptive immune phenotypes. In parallel, activation-associated programs were broadly enriched in T-cell populations, collectively suggesting that statin treatment not only increases T-cell abundance but also promotes adaptive immune remodeling. Notably, CD8^+^ T-cell populations appeared to undergo more pronounced remodeling compared with CD4^+^ T cells. These findings are further supported by our ex vivo studies, in which we found that CD8^+^ T-cell populations exhibit greater phenotypic responsiveness to ATO-conditioned ascites, with increased expression of IL-2 and IL-15Rβ, compared with CD4^+^ T cells. Together, these data indicate that the daily clinical dose of statins preferentially promotes adaptive CD8^+^ T-cell engagement in the tumor microenvironment.

In both the in vivo and ex vivo analyses, increased PD-1 expression was observed in T-cell populations after ATO treatment. Traditionally, PD-1 has been regarded as a canonical immune checkpoint molecule associated with T-cell exhaustion and tumor immune evasion. However, this interpretation appears inconsistent with the concurrent T-cell activation observed in our study, including enrichment of the effector CD8^+^ T-cell program, and TLS activation and maturation. One possible explanation is that T cells within the statin-remodeled tumor microenvironment may adopt complex and dynamically regulated phenotypic states, in which activation-associated and checkpoint-associated features coexist, as shown in a recent study (16). Increasing evidence suggests that statin treatment may enhance anti-PD-1 therapeutic responses in certain tumor settings, including melanoma and non-small cell lung cancer (17–19). Therefore, the observed PD-1 upregulation may reflect complex immune remodeling induced by statin use, which creates an opportunity for combination treatment with immune checkpoint blockade.

Macrophages represent one of the major immune cell populations in malignant ascites. We did not observe a significant change in the ratio of pro-inflammatory M1 and pro-tumorigenic M2 cells in ATO-versus DMSO-treated mice. In scRNA-seq analysis, we observed that ATO treatment may enhance IL-15 expression in the ascites macrophages and myeloid cells. Consistently, ex vivo experiments confirmed that ATO increased IL-15 expression in BMDMs both directly and indirectly through tumor supernatant. IL-15 is a cytokine that supports CD8^+^ T-cell survival, activation, and effector differentiation under inflammatory conditions. In light of these findings, we propose that IL-15 stimulated by ATO-primed macrophages may play an important role in enhancing T-cell immunity, thereby contributing to the improved overall survival.

Overall, our findings in an immunocompetent murine model indicate that clinically relevant low-dose statin use reshapes the tumor immune microenvironment through coordinated remodeling and enhances overall T-cell immunity. The findings suggest the underlying mechanism of the prolonged survival of statin-using cancer patients. Statins are among the most widely used and prescribed medications in the world. Globally, more than 200 million adults take statins regularly to reduce blood cholesterol and prevent associated cardiovascular diseases. In the United States, approximately 40-92 million adults take statins. Among the various statins, atorvastatin (brand name Lipitor) is the most widely used, accounting for 36% of prescriptions, followed by simvastatin at 34%. Both of these statins are lipophilic and dissolve easily in lipid-rich media (20). The current study provides evidence that even after a cancer diagnosis, there is no evidence of harm associated with continued statin use. Instead, there is even a potential survival benefit. Such benefits of statin use may extend beyond ovarian cancer patients, as studies have shown that prolonged statin use is linked to reduced cancer-associated mortality across multiple cancer types, including endometrial, blood, breast, lung, and gastrointestinal cancers (8,21,22). Therefore, findings from the current study provide a basis supporting the potential of a low-cost, simple, and relatively safe approach to the clinical management of cancer patients. This suggests that statins could serve as a broadly applicable and accessible adjunct in cancer care.

## Declarations

### Availability of data and materials

The datasets used and/or analyzed during the current study are available from the corresponding author upon reasonable request.

### Competing interests

The authors declare that they have no competing interests.

### Funding

The study was supported by NIH/NCI P50CA228991, NIHR01CA260628, DoD W81XWH-22-1-0852, and Tina’s Wish Team Award.

### Author contributions

JML and SQ designed and conceptualized experiments and methodology, performed experiments, analyzed data, and wrote and edited the manuscript. MR developed the multiplex staining protocol and carried out related experiments. TLW & IMS provided conceptual inputs, offered critical insights, wrote and edited the manuscript.

## Acknowledgments

We thank Dr. John Tobias at the University of Pennsylvania for assistance with the processing and integration of the single-cell RNA sequencing data, including Seurat-based clustering and cell-type annotation. We also thank Xiaoling Zhang of the Johns Hopkins Ross Flow Cytometry Core for her technical assistance with FACS sorting and the MGLFRC lab members for kind technical assistance and experimental help.

## Supplemental information

Figures S1-9

Supplementary Table

Supplementary Materials and Methods

Supplementary File: 3D Umap.mov

STAR Table

## Notes

### Competing Interest Statement

The authors have declared no competing interest.

